# An Equivariant Bayesian Convolutional Network predicts recombination hotspots and accurately resolves binding motifs

**DOI:** 10.1101/351254

**Authors:** Richard Brown, Gerton Lunter

## Abstract

**Motivation:** Convolutional neural networks (CNNs) have been trememdously successful in many contexts, particularly where training data is abundant and signal-to-noise ratios are large. However, when predicting noisily observed biological phenotypes from DNA sequence, each training instance is only weakly informative, and the amount of training data is often fundamentally limited, emphasizing the need for methods that make optimal use of training data and any structure inherent in the model.

**Results:** Here we show how to combine equivariant networks, a general mathematical framework for handling exact symmetries in CNNs, with Bayesian dropout, a version of MC dropout suggested by a reinterpretation of dropout as a variational Bayesian approximation, to develop a model that exhibits exact reverse-complement symmetry and is more resistant to overtraining. We find that this model has increased power and generalizability, resulting in significantly better predictive accuracy compared to standard CNN implementations and state-of-art deep-learning-based motif finders. We use our network to predict recombination hotspots from sequence, and identify high-resolution binding motifs for the recombination-initiation protein PRDM9, which were recently validated by high-resolution assays. The network achieves a predictive accuracy comparable to that attainable by a direct assay of the H3K4me3 histone mark, a proxy for PRDM9 binding.

**Availability:** https://github.com/luntergroup/EquivariantNetworks

**Supplementary information:** Supplementary data are available at *Bioinformatics* online.

## 1 Introduction

Deep Learning based models have been highly successful in many areas where traditional modeling approaches appeared to have reached their limits. This is true also for modeling biology from sequence, where Deep Learning sequence models have been shown to outperform previous state of the art techniques (Zhou and Troyanskaya, 2015; Alipanahi *et al.*, 2015; Kelley *et al.*, 2016). These models have several attractive characteristics, including their ability to learn without the need for manual feature curation or model seeding, and the ability to learn complex nonlinear interactions. This is balanced by the need for large amounts of training data, and the tendency of these models to overtrain. In many situations this is not problematic, but for applications in biology training data is often fundamentally limited, either by the size of the genome, the limited genetic diversity of a population, or the cost of assaying individuals or samples. In this context it is particularly important to exploit the known structure of the model as much as possible, and to try and avoid overtraining and improve generalizability.

As an example of a biologically motivated problem with limited training data, we consider the problem of predicting recombination hotspots from sequence. In humans, the rate of meiotic recombination varies greatly along the genome, with recombinations occurring primarily in short regions colloquially known as recombination hotspots. The mechanism for this localisation has been shown to be the action of the zinc finger protein PRDM9 (Baudat *et al.*, 2013). After being expressed in meiotic prophase, PRDM9 binds DNA in a sequence-specific manner, and catalyzes H3K4 and H3K36 trimethylation and double stranded breaks, some of which are resolved as recombinations.

The canonical PRDM9 binding motif, CCTCCCTNNCCAC, was identified by an enrichment analysis of sequences underlying hotspot vs. those in regions not involved in recombination (Myers *et al.*, 2008). However, while significantly enriched, this motif is only weakly predictive of recombination. For example, in our data it appears in around 2% of hotspots and 0.3% of coldspots. This, coupled with the fact that there are only around 20,000 hotspots, many of which are ill resolved (median length approximately 2000 base pairs (bp)), makes prediction of recombination hotspots challenging.

To build a model that optimizes predictive power given these constraints, we combine two recently introduced ideas. One is that of equivariant convolutional networks (Cohen and Welling, 2016) which we use to build a network exhibiting the reverse-complement symmetry of double-stranded DNA. Equivariance is a richer concept than invariance; while a sequence and its reverse-complement are expected to exhibit the same predisposition for recombination, the binding of proteins to DNA is usually not reverse-complement symmetric, and protein-protein interactions can be similarly directional. This is reflected by symmetries on higher levels in the convolutional neural network (CNN) that mirror the reverse-complement symmetry on the sequence level. Equivariance is the mathematical concept that describes how the action of reverse-complementing is reflected in the different layers of the model.

The second insight is that dropout, a commonly used regularization technique for CNNs, can be interpreted as an approximation of a variational Bayesian inference (Gal and Ghahramani, 2016), here referred to as Bayesian dropout. This interpretation suggests particular modifications of standard applications of dropout. In particular, this interpretation suggests the use of Monte Carlo averaging of activations (MC dropout) rather than weight averaging at the prediction stage. The implementation suggested in the literature would break equivariance (Gal and Ghahramani, 2016). We here show how to obtain an equivariant version of Bayesian dropout, and how to use this to obtain a model exhibiting exact reverse-complement symmetry while retaining the advantages of Bayesian dropout.

The remainder of the paper is organized as follows. In section 2 we introduce equivariant networks, and show how to build equivariance into standard CNN layers including convolutional layers, max-pooling and dropout. We show how to make the Bayesian dropout scheme equivariant, and we introduce a new max-pooling layer that acts over the action of the reverse-complement (RC) symmetry group. In section 3 we show that RC-equivariant networks and Bayesian dropout each and in combination significantly increase predictive accuracy, both on simulated and real data, whereas classical dropout hurts performance. We further show that our network outperforms state-of-art motif finders, and is able to identify high-resolution binding motifs. We finish with discussion and conclusions in Section 4.

## 2 Methods

### 2.1 Equivariant networks

We use feedforward neural networks to generate a learnable mapping directly from the input sequence to an output response variable, which in this case is a class assignment. The network is composed of a sequence of layers that define a directed acyclic computation graph, with each layer acting in turn on the output of its ancestor. Explicitly, for a 2 dimensional input tensor *X*_*ij*_, the network can be viewed as a function composition

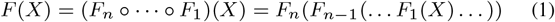

with component functions *F*_*i*_ representing the actions of layer *i*; here ○ denotes function composition. Note that component functions are defined on their own tensor spaces, *F*_*i*_:*T*^(*i*−1)^ → *T*^(*i*)^.

In our case, *X*_*ij*_ represents a length-*N* DNA sequence from an alphabet {*A*,*C*,*G*,*T*} which is usually one-hot encoded (Lanchantin *et al.*, 2016; Alipanahi *et al.*, 2015; Zhou and Troyanskaya, 2015) so that *T*^(0)^ = ℝ^4×*N*^. The DNA represented by the input sequences *X* exists physically mostly in a double-stranded form, with one strand hydrogen bonded to its reverse-complement. This means that the sequence seen by the model could just as naturally be represented by its reverse complement, and the network should arrive at identical outputs for these two sequences:

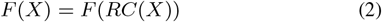

where *RC*:*T*^(0)^ → *T*^(0)^ maps the encoding of a sequence to the encoding of its reverse complement. One way to achieve this symmetry is to require that *F*_1_(*RC*(*X*)) = *F*_1_(*X*). However, this is highly restrictive; the first layer often represents protein binding motifs, which are often not RC symmetric. Intuitively, one wants the output of a layer to exhibit the “equivalent” symmetry appropriate for the encoding of the next layer. The mathematical translation of this is to require equivariance:

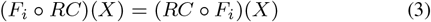

for all *i* and all *X* ∈ *T*^(*i*−1)^. A graphical representation of this relation is given in Figure 1. Note that the two operators *RC* in equation (3) are different, as they act on different tensor spaces. In particular, in our case *RC* acts as the identity on *T*^(*n*)^ to ensure that the full model is invariant under *RC*, and on *T*^(0)^ the operator *RC* is determined by the chosen encoding. The modeler has freedom in choosing *RC* on intermediate layers, subject to the constraint (3), which also imposes constraints on the *F*_*i*_, the initialisation, training procedure, and any parameter ties used during training. Note that this setup is not restricted to RC equivariance, and is valid for any group with actions on tensor spaces, including mirror symmetries, rotations and translations. In this general case the operators *RC* in Figure 1 are replaced by group actions 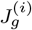 operating on *T*^(*i*)^, where *g* is a group element. We will not pursue that direction here, but see Cohen and Welling (2016) for an exposition.

**Fig. 1.**
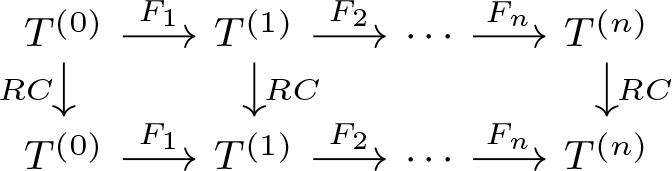
Commutative diagramfor areverse complementequivariantnetwork. Compositions of functions along any path in the network depend only on the start and end point, and not on the path taken.

### 2.2 Choice of one-hot basis

We define the vectors

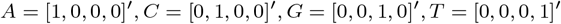

where ′ denotes transposition, and encode a genomic sequence by the concatenation of corresponding column vectors. For a sequence encoded in this way,

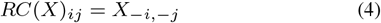

with negative indices denoting offsetting from the opposite end of each dimension by that amount.

### 2.3 Convolutional layer

For sequence classification we use a 1D convolution layer, with *n*_*f*_ filters of length *f*_*l*_ stored in a weight matrix *W*_*ijk*_. The output of such an operation on tensor *X*_*ij*_ (ignoring bias terms for simplicity) is

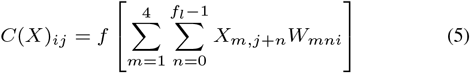

where *f* is the activation function. Note that we used “valid padding” so the sequence dimension is reduced by *f*_*l*_ − 1. Applying the reverse complement operation to the input tensor of shape [4, *N*] yields

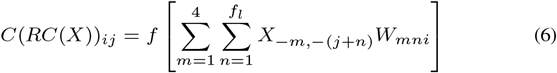

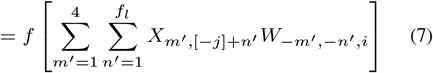

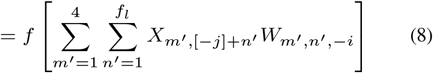

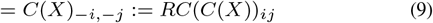

where we used the substitutions *m* = −*m*′, *n* = −*n*′ = *f*_*l*_ + 1 − *n*′, and [−*j*] denotes the positive index *N* + 2 − *f*_*l*_ − *j*. At (8) we assumed that *W* obeys the symmetry *W*_*m,n,i*_ = *W*_−*m*,−*n*,−*i*_. Therefore, the convolutional layer satisfies (3) if this weight symmetry holds, and if we define RC on the output layer as in (9). A schematic of this is shown in figure 2A. We note that this specific symmetry was used in a convolutional layer in Shrikumar et al. (2017) and was shown to improve inference.

### 2.4 Max-pool layer

Spatial Max Pooling along the position dimension of a tensor is used in order to increase the receptive field at the expense of resolution. We consider the special, commonly used, case where the stride length is the same as pool width. For this case we have for a pool length *p*_*l*_,

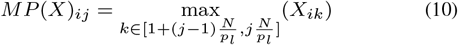

As long as *p*_*l*_ divides *N* this defines an equivariant mapping, with *RC* defined on the next layer in the obvious way.

### 2.5 Reverse complement max pooling

We frequently found it useful to pool along the “filter axis” rather than along the spatial direction. More precisely, we take maxima along orbits under the group action, which in our case consist of two elements. After RC max pooling we therefore have a new output

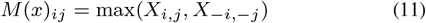

This process halves the size of the output tensor compared to the input. The resulting compression is depicted in figure 2B. Equation (3) is again satisfied, now with *RC* acting as the identity on the new layer. A network that contains RC max pooling is therefore automatically symmetric under reverse complementing. In general, more complex symmetry groups may contain nontrivial subgroups, and any of those may be used to do partial symmetric max pooling.

**Fig. 2.**
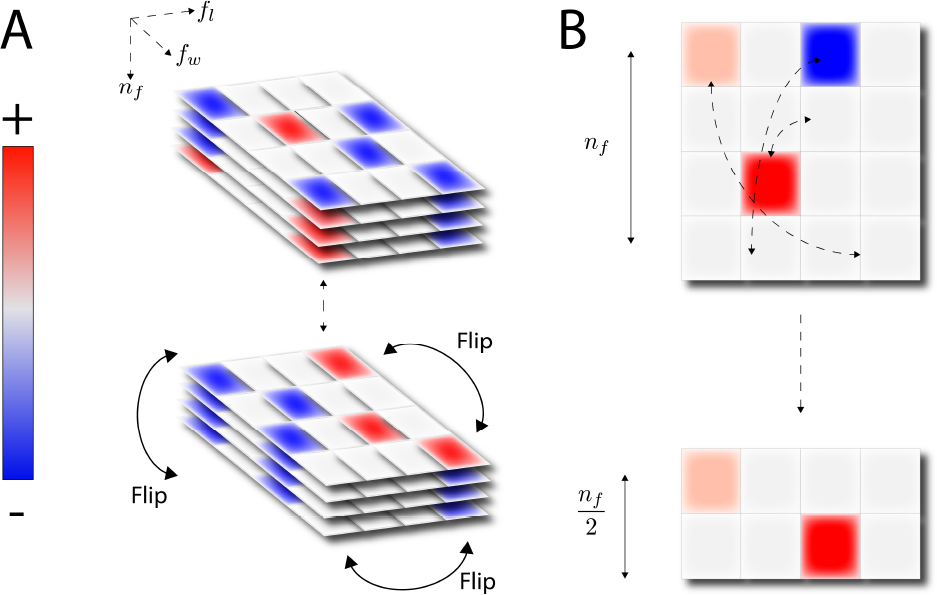
Left: The 3 dimensional filter tensor has enforced symmetry by weight tying by flipping the second half of the filter axis as shown. Right: An contrast to a conventional max pooling along the spatial dimension, this approach permits pooling along filters.

### 2.6 Dropout layer

Dropout is a form of stochastic regularization for neural networks designed to prevent over-fitting by adding noise to the output of a layer during network training (Srivastava et *al.,* 2014). This is commonly implemented by dropping out nodes according to a Bernoulli-distributed random variable, and can be implemented by taking the Hadamard product between two identically shaped tensors. Define

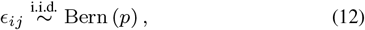

then dropout applied to a tensor *X* with dropout rate *p* is

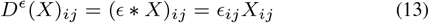

with the tensor *ϵ* sampled on a batch-wise basis. We denote the Hadamard product by * rather than the more usual ○ to avoid confusion with function composition. Applying *RC* gives

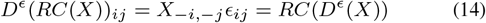

which holds if

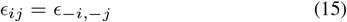

and if *RC* acts in the same way on the output as on the input layer.

### 2.7 Bayesian Equivariant networks

Convolutional networks have been shown to work well on large datasets, but it is known that they overfit quickly when relatively little training data is available (Gal and Ghahramani, 2016). This is often the case in the biological domain, calling for a principled approach to deal with limited training data sets. Networks with dropout after every convolutional layer and trained via backpropagation can be seen as approximating Bayesian variational inference (Gal and Ghahramani, 2016), promising good behaviour in data-poor settings. The interpretation also suggested to use Bayesian dropout rather than traditional weight averaging for making predictions, using dropout to approximate sampling from a posterior weight distribution. In this interpretation, the corresponding prediction is obtained by the average over a sample of instantiations of the network:

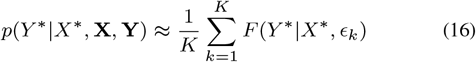

for unseen *X* * given previous training examples **X** = (*X*_1_,…, *X*_*n*_) and **Y** = (*Y*_1_,…,*Y*_*n*_) used to train *F*. The set of random variables ϵ = (ϵ_1_,…,ϵ_*K*_) implement a random sample of the weights defining the network *F*, and as long as these obey the correct symmetry, e.g. (15), each term will be RC-symmetric and so will the sum. The resulting function will be stochastic, but as long as ϵ is sampled and fixed beforehand, it will nevertheless exhibit exact RC symmetry.

### 2.8 Choice of activation function

Rectified linear units (ReLUs), defind as ReLU(*z*) = max(0,*z*), are activation functions applied at the terminus of every layer, providing the requisite nonlinearity for stacked layers to have richer representational power than a simple linear model. These activation functions have become ubiquitous in deep learning, though usually for networks which are composed of more layers than is typical in genomic problems, as they reduce or resolve the vanishing gradient problem. ReLU elements have the property that for a large number of units, the output will be identically zero. Although sparsity can be advantageous, e.g. by making interpretation easier, we found that it hampers convergence in all problems we have tried. We found thatusingELUs (Clevert *et al.,* 2015)resultin substantially better convergence behaviour on our dataset (see Supplementary Fig. 1). We observed qualitatively similar results using shifted ReLUs (SReLU(*z*) = max(*z*, − 1)) (data not shown).

### 2.9 Initialization of the output layer

We found that using a custom initialization of the output layer, providing the classification scores before a final softmax transformation, substantially improved convergence for the networks we considered (Supplementary Fig. 2). We initialized the weight matrix of the final layer with 1, and the two bias parameters (corresponding to the two nodes representing class probability) with {1, −1}. A motivation for this choice was the observation that on a number of problems the output layers weights were always close to these values, so it appears that this initialization is closer to the global optimum than traditional approaches such as Glorot and Bengio (2010).

### 2.10 Network architecture

We performed a two separate hyperparameter searches, one for the simulated data and one for the recombination dataset. For the recombination dataset, we varied the number of convolutional layers (*n* ∈ {1, 2,…5}), the filter length of the input layer (*f*_*l*_ ∈ {10, 15, 20, 30}), the filter lengths of each internal convolutional layer 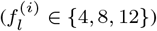, the max pooling layer sizes after each convolutional layer (*p*_*l*_ ∈ {4,8,12}) and the number of filters at at each convolutional layer 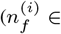 {8,16,32,64}). In additionwe optionally added an L^2^ regularization term of varying strength (*α* ∈ {0, 0.0001,0.0003, 0.001}). For the simulated dataset we knew there were no interactions, and we knew the length of the motifs of interest, so we used a single convolutional layer and optimized the number of filters (*n*_*f*_ ∈ {4, 6, 12}), along with the L^2^ penalty term. The configuration of the final networks is provided in Supplementary Material section A.

## 3 Results

### 3.1 Datasets

Simulated data was generated as follows. As a model for a regulatory network involving two binding proteins, we sampled two PWMs (ATAF4 and ERF1) from JASPAR (Sandelin *et al.*, 2004). We first randomly sampled 40,000 times a random {0, 1}-response with 50% probability for each, representing a measured phenotype of interest. For each response variable with value 1 we sampled a specific motif from each of the two PWMs (reverse complementing them at random) and injected it into a random background sequence with 40% probability. Otherwise we injected the motifs with probability 20%. This procedure resulted in 40000 sequences of length 1000, with 20110 in category 1 and 19890 in category 0. Note that there is a substantial amount of noise in this dataset, making for a challenging classification problem.

We also prepared a dataset of human recombination hot-and coldspots, similar to Myers *et al.* (2008). We first applied a simple hidden Markov model to segment the genome into regions classified as ‘hot’ and ‘not hot’. For emission probabilities we used two exponential distributions for *p*(observed rate|hot) and *p*(observedrate|nothot), and used Viterbi training to set the parameters. The median recombination rate in regions classified as ‘hot’ was 10.5 cM/Mb. The length of hotspots, once annotated, had a median value of 2448 bp, but with a heavy tail up to around 20 kbp. For our purposes, we wanted to study localized recombination events, so we discarded all hotspots longer than 4 kb. After discarding sequences with unspecified bases, we then sampled a sequence of length 1kb from the centre of the remaining hotspots, yielding a total of 17552 truncated hotspots for classification.

To define a matched set of coldspot regions once hotspot regions were identified, we applied a greedy search within 300 kb of each hotspot region to identify a sequence within 10% GC content, and with a recombination rate below 0.5 cM/Mb. GC matching ensures that GC content, which tends to be higher in recombination hotspots due to GC-biased gene conversion, cannot be used as a proxy for recombination strength, forcing the network to focus on causal signals of DNA binding motifs. This pipeline yielded a total of 17547 truncated coldspots, also of length 1000 bp.

### 3.2 ELU ans SReLU activations improve convergence

Across a wide range of learning and topology hyperparameters we observed consistent difficulty with the ReLU activation function, with networks often not converging to optimal accuracies, and sometimes not converging at all, leaving no better than random guesses. We observed, upon experimentation, that ELU and SReLU activation functions, gave much improved results (see Supplementary Figure 1 for an illustration of this effect with ELU activations).

### 3.3 Equivariant Bayesian networks improve classification accuracy

We first identified two optimal non-equivariant networks by performing a hyperparameter search across filter number and lengths, max pool layers, regularization parameter and number of layers as described above, independently for the simulation data and recombination data. After optimization, we then considered reverse-complement equivariant networks, and re-optimized the number of kernels and regularization parameter for both data sets independently, keeping all other parameters fixed. Orthogonally to this, we also added Bayesian dropout to the baseline networks (removing L^2^ regularization if it improved validation accuracy) and added equivariant Bayesian dropout to the equivariant networks, yielding four independently optimized models.

Comparing the best-performing network in each of the 4 categories we found that for both datasets the equivariant Bayesian networks outperformed the best-in-class non-equivariant networks, achieving significantly better test accuracy over a sample of 50 runs (Figure 3). We found that both equivariant non-Bayesian and Bayesian non-equivariant networks were significantly more accurate than the best in class convolutional network, and that the combination of Bayesian dropout and equivariance was again significantly more accurate than either.

We also investigated how well the equivariant Bayesian network performed for these two problems in comparison to classical data augmentation, whereby the reverse complement of every sequence is added to the dataset. Although data augmentation did improve the accuracy over standard networks trained without data augmentation, we found that the equivariant Bayesian networks significantly outperformed classical network even when trained with an augmented dataset, for both problems (Supplementary Fig. 4).

**Fig. 3.**
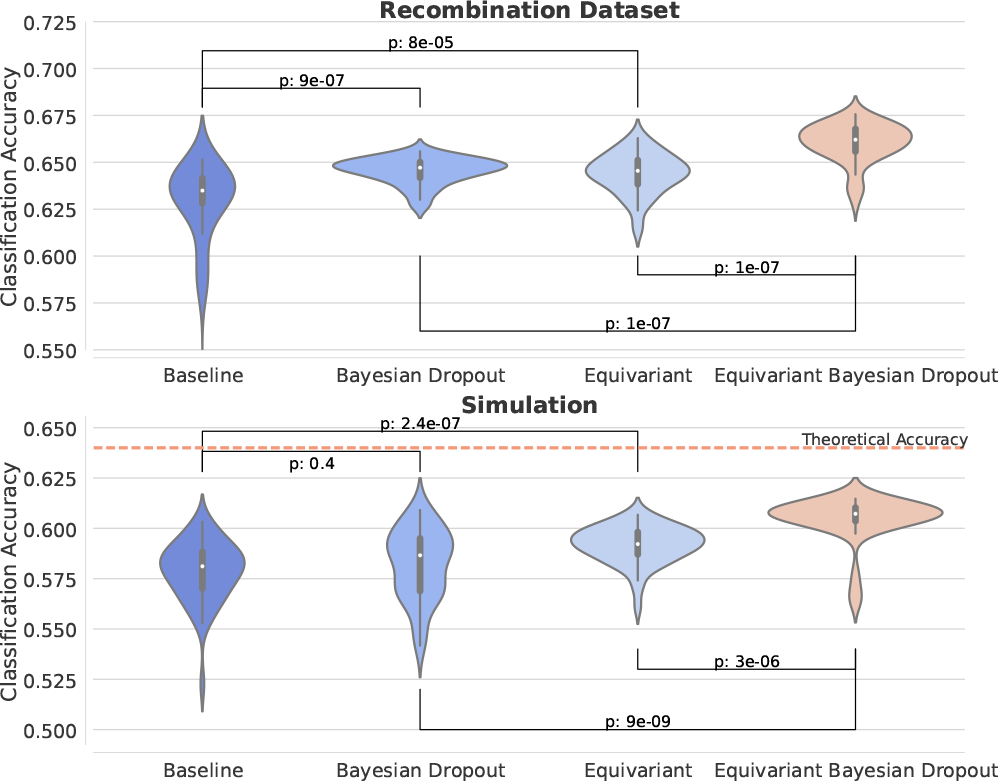
For every network configuration, we trained to convergence a total of 50 times and built distribution of the final test accuracy. Note the for both the simulation (top) and the recombination data (bottom) we find a statistically significant improvement over the case where this symmetry is not applied

### 3.4 Conventional dropout considered harmful

To understand the contribution of Bayesian dropout on the performance of the network, we compared the final equivariant topology for both datasets with no dropout, Bayesian dropout and conventional dropout, recording the mean accuracy over 50 trials. Compared to Bayesian dropout, an identical network that used classic dropout procedure yielded substantially inferior results (table 1), and performs worse than the baseline without dropout. This behaviour was seen in both datasets.

**Table 1.** Mean accuracies and errors for the symmetric networks with no dropout, Bayesian dropout and conventional dropout

| Dataset | Baseline | Bayesian Drop | Classic Drop |
| --- | --- | --- | --- |
| Sim | 59.2 ± 0.1 | **60.4 ± 0.2** | 56.0 ± 0.2 |
| Recomb. | 64.4 ± 0.2 | **66.0 ± 0.2** | 56.4 ± 0.3 |

### 3.5 Equivariant Bayesian networks improve upon existing motif finders

To assess our approach we compared our results with the state-of-art neural-network-based motif finder DeepMotif (DeMo; Lanchantin *et al.* (2016)) on the recombination dataset. We benchmarked our results against the three network configurations offered by the package, CNN, RNN and CNNRNN. All of these have substantially more complexity and weights than the model we used, but failed to perform well on this task. Indeed the CNN model (a convolutional neural network) often failed to converge at all, a problem that we also observed when training our vanilla convolutional networks on these datasets. The first training attempt that converged to better than random predictions had AUROC 0.58. Both the RNN and CNNRNN toplogies converged reliably, but achieved AUROCs of 0.64, substantially less than the median AUROC of 0.71 achieved by our equivariant network. We note that the CNNRNN model appeared to overfit slightly and may have benefited from an early stopping regime in the original code. Nonetheless, the peak validation accuracy was only marginally improved, and not close to the accuracy of the Bayesian equivariant model.

**Fig. 4.**
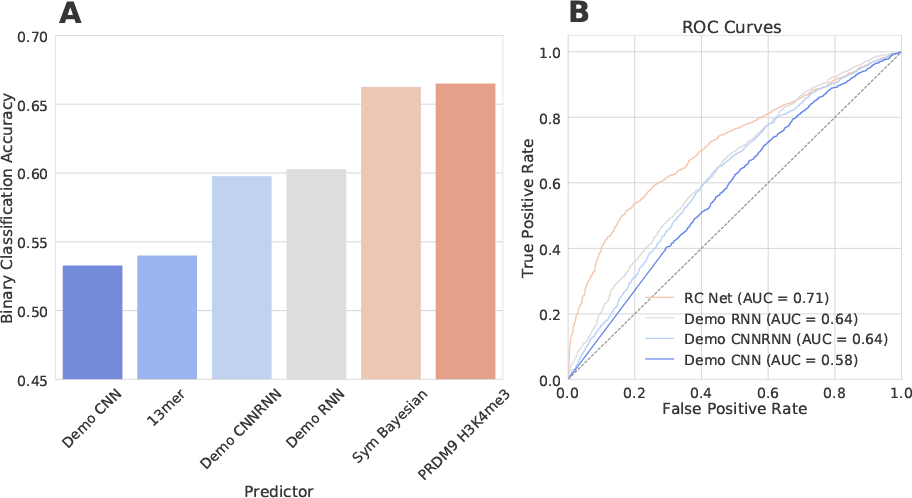
Classification accuracy (left) and (area under) receiver operator curves (right) for the equivariant Bayesian network and the three models provided by the Deep Learning based motif finder DeMo (Lanchantin et al., 2016). For comparison we also show the classification accuracy of the canonical PRDM9 13mer motif, as well as the classification accuracy obtainable by using the differential histone mark H3K4me3 under PRDM9 overexpression, a direct proxy for PRDM9 binding (Altemose et al., 2017).

### 3.6 Discovery of novel PRDM9 binding motifs

Using the equivariant Bayesian network with the best classification accuracy, we identified binding motifs by identifying a subset of input sequences responsible for the activation of particular nodes at the first layer, the subsequence driving the activation in each of those, and building a Position Weight Matrix (PWM) from the aligned subsequences.

This process identified five motifs, three of which are versions of the classical 13mer that was identified via enrichment analysis and exhaustive search of motifs in Myers *et al.* (2008) (Fig. 5c-e). In addition we identified two substantially longer (22 nt) and sparse motifs that were not previously identified using this data set (Fig. 5a,b). Since our networks are only approximately Bayesian, to confirm that these motifs were not artefacts of overtraining, we confirmed their significance by frequentist tests of significance for enrichment on hold-out data (Table 2). The enrichment of the sparse motif is similar to that of the canonical 13mer originally associated with PRDM9 binding, but its complexity and under-constrained nature (only about 8 out of the 22 bases of the motif have substantial information content) mean that it was not originally discoverable using a traditional enrichment approach.

## 4 Discussion

Deep learning approaches have been shown to be very effective in building sequence models on large scale data (Zhou and Troyanskaya, 2015; Kelley *et al.,* 2016; Quang and Xie, 2016). However, through simulated and biological data we show here that models designed using traditional building blocks for neural networks may struggle to converge consistently and produce reliable results in cases where the signal in the data is weak, and the amount of training data is limited. This problem was seen across a large number of network architectures, as well as in methods specifically designed to identify transcription factor binding sites from sequence data (Lanchantin *et al.,* 2016).

**Table 2.**
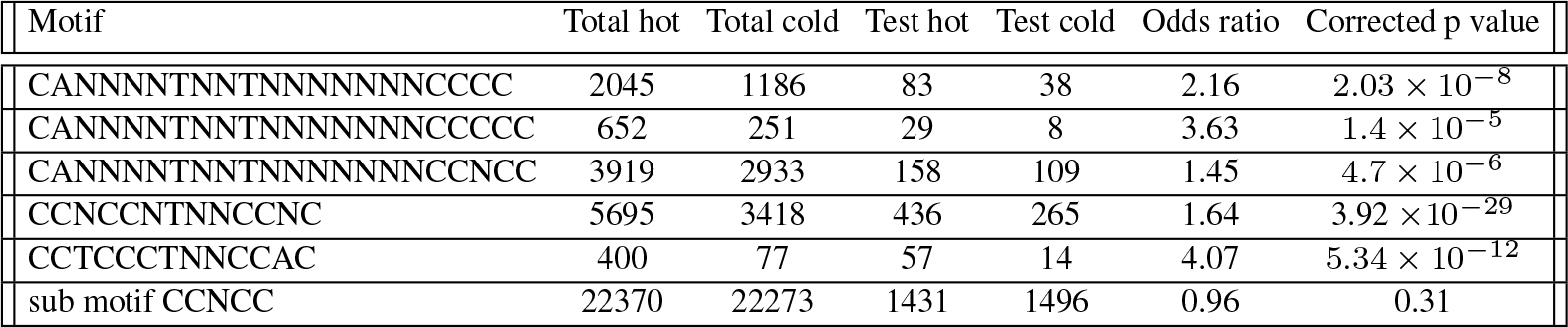
Comparison of the novel motif with previously known PRDM9 binding site in both the training and test datasets. Total hot/cold, number of motifs in the full hotspot/coldspot data set; test hot/cold, number of motifs among hold-out test data consisting of 1454 hotspots and 1530 coldspots; corrected p value, Bonferroni-corrected p values for association (Fisher’s exact test)

**Fig. 5.**
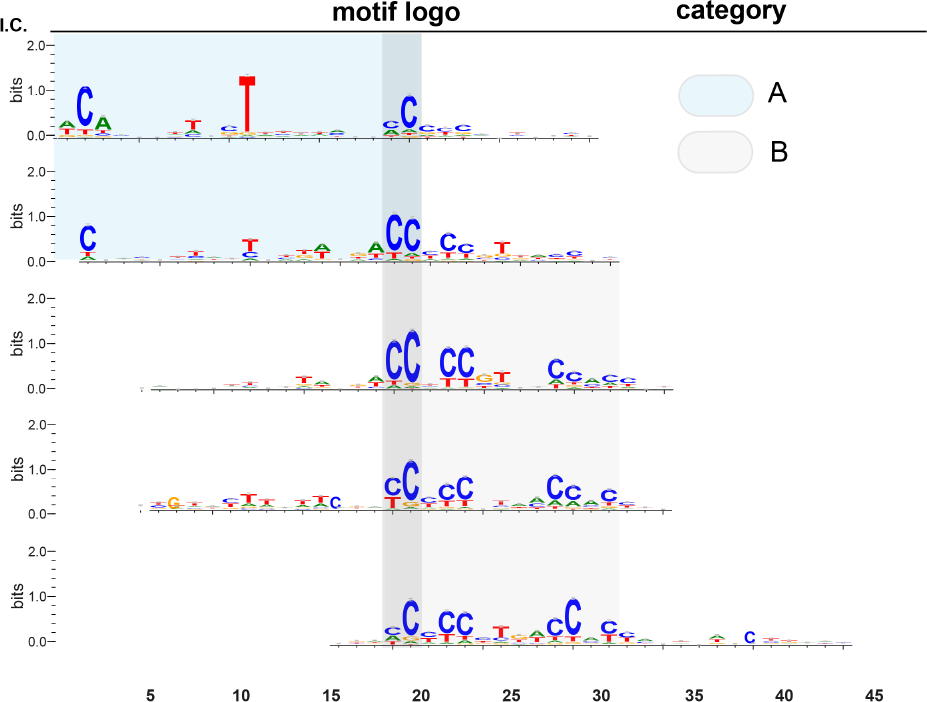
Top: We found the sequences corresponding to the maximum activation on each filter, and from these built sequence logos. We found a number of motifs (category B) corresponding to the classical PRDM9 binding motif that was discovered with an exhaustive search of an analogous dataset. Additionally, we found 2 novel 22base motifs, that were previously undiscovered on this dataset. It is notable that they bear a striking resemblance to the recently discovered motif by Altemose et al. (2017).

To improve on this situation, we showed how to combine equivariant neural networks (here, neural networks that exhibit exact reverse-complement symmetry) with Bayesian dropout. While a naive combination of these ideas would result in networks that are only in expectation reverse-complement symmetric, we show that it is possible to achieve exact RC symmetry. In addition, we find that by modifying the activation functions, and the initialization of the output layer, we obtain a further significant improvement in accuracy.

Equivariant networks can be implemented in several ways. We chose to implement a standard (non-equivariant) network to and enforce equivariance by requiring certain identities on the parameters. Although this introduces some extra computation, this approach compares favourably with data augmentation, an alternative approach to handling symmetries where the training data is made symmetric, and symmetry must be learned by the network. Data augmentation applied to RC symmetry doubles the training time, and we show that although this data augmentation improves results over baseline models, an equivariant network achieves significantly better results at lower computational cost.

We found that Bayesian dropout also resulted in a substantial improvement in the performance. This is striking, as dropout is normally though of as a regularization technique that reduces the tendency of overtraining often found in large model. By contrast, in our regime the final models were small in comparison to the number of samples in the training set, and we did not see much evidence of overfitting either with or without using dropout: we did not see substantial continued decrease of training loss after test loss stabilised, and we did not find that the model latched on to spurious motifs that were not statistically significant on test data. It appears that, in addition to addressing overtraining, Bayesian dropout leads to superior learning and feature extraction. It would be interesting to confirm this phenomenon in different settings.

We then applied this methodology to predict meotic recombination hotpots directly from sequence. It was straightforward to interpret the resulting model, and interrogation of the sequences that maximally activated the input layer revealed the previously characterized 13 base PRDM9 binding motif (Myers *et al.*, 2008). Additionally, we discovered a much sparser motif, with only about 8/22 bases showing substantial information content. Like the classical motif, these new motifs were statistically significant on hold-out data, indicating that they were truly predictive features and not artifacts of an overtrained model. These sparse motifs (category A in figure 5) bear strong resemblance to motifs recently shown to be associated with PRDM9 binding Altemose *et al.* (2017), lending further support to this conclusion. In fact, the observation, presented in figure 4, that the model was able to achieve predictive accuracy on a par with hotspots predicted using differential H3K4 trimethylation as an input (also from Altemose *et al.* (2017)) is consistent with the hypothesis that the neural network model represents a near-optimal model for PRDM9 binding. The alternative explanation is that our model and the the H3K4me3 assay are suboptimal to the same degree.

From a practical point of view, we noted that the motifs generated from this network with Bayesian dropout were qualitatively different and more numerous than the motifs identified with a classical convolutional approach, and we see a degree of degeneracy among the motifs learned by our network. It remains to be seen whether these different binding motifs correspond to related but slightly different binding modalities, or whether these motifs are a result of dropout training and provide a way for the network to robustly identify motifs in the presence of weight noise. This would be an interesting direction fur further research. Certainly, if indeed these motifs do correspond to different binding modalities, each with slightly different binding affinities, it would explain why these networks are able to achieve the superior classification accuracies compared to standard models.

## Acknowledgements

We thank Simon Myers for helpful discussions.

